# In Silico Modelling of Shear Stress and Energy Dissipation Rate Effects on Human Pluripotent Stem Cell Proliferation in Vertical-Wheel Bioreactors

**DOI:** 10.64898/2026.04.22.720266

**Authors:** Ferdinand R. Avikpe, Faisal J. Alibhai, David A. Romero, Amirmahdi Mostofinejad, Julia E.S. Bauer, Coulter Montague, Michael A. Laflamme, Cristina H. Amon

## Abstract

Human pluripotent stem cells (hPSCs) hold significant promise for regenerative medicine, yet optimizing their expansion in three-dimensional bioreactor systems remains challenging due to complex interactions between mechanical forces, metabolic constraints, and aggregate formation dynamics. This study developed and validated a mechanistic mathematical model to predict hPSC proliferation dynamics in vertical-wheel bioreactor (VWBR) systems, incorporating the effects of shear stress and energy dissipation rate (EDR) on cell growth and aggregate dynamics. Seven model variants employing different kinetic formulations for shear stress and energy dissipation rate effects were systematically evaluated through model selection, identifiability analyses, and experimental validation. Experimental data from six bioreactor conditions varying in initial cell density (2 × 10^4^–15 × 10^4^ cells/mL), agitation rate (30–60 RPM), and working volume (100–500 mL) were used for model calibration and selection. Bayesian Information Criterion analysis identified a model combining Michaelis-Menten kinetics for shear stress inhibition with a EDR-mediated aggregate detachment formulation as the best-performing variant, achieving a Mean Relative Prediction Error of 13.97%, comparable to the experimental variability of 16.29%. Independent validation experiments using leave-out data gathered under different media exchange schedules confirmed model accuracy with prediction errors below 14%, consistent with observed experimental variability around 12%. The validated model was used to optimize the media exchange protocol, leading to a 37.5% reduction in media consumption with only a 13.5% reduction in final cell yield, demonstrating its utility for prospective, quantitative bioprocess design in VWBR systems.

## 1 Introduction

Human pluripotent stem cells (hPSCs), including embryonic (hESCs) and induced (hiPSCs) pluripotent stem cells [1, 2], have emerged as a cornerstone in regenerative medicine and tissue engineering due to their ability to self-renew and differentiate into multiple cell types of the human body [3, 4, 5]. These properties make hPSCs invaluable for applications ranging from disease modelling and drug screening to cell-based therapies [6, 7, 8, 9]. Despite advancements in bioreactor technology and culture media formulation, optimizing hPSC culture conditions in 3D as aggregates remains a fundamental challenge, particularly in the context of expansion/proliferation dynamics. The ability to expand hPSCs *in vitro* while preserving their pluripotency is highly dependent on a range of biochemical and mechanical factors, including nutrient availability, cell-cell interactions, and environmental stresses [10, 11, 12].

*In silico* modelling, the use of mathematical models and computational tools to simulate biological processes, is becoming an essential tool for understanding complex biological systems, and hPSC culture is no exception [13]. Accurate models of hPSC growth dynamics can serve as predictive tools to optimize culture conditions, improve scalability, and reduce experimental costs [14, 15]. By integrating experimental data with computational frameworks, these models facilitate rational decision-making in bioprocess design and provide mechanistic insights into cell behaviour under different culture conditions. Crucially, the predictive power of in silico analysis helps constrain the parameter space during scale-up processes, thereby limiting the substantial costs associated with empirical protocol optimization through extensive trial-and-error experimentation.

Several mathematical models have been employed to study cell growth across various cell types, such as neural cells [16, 17, 18], red blood cells [19], bone [20, 21, 22, 23, 24] and cartilage [25, 26], among others. These models generally fall into two categories: ordinary differential equation (ODE) models, which describe how cell populations and biochemical interactions evolve over time in well-mixed conditions, and Advection-Diffusion-Reaction (ADR) models, which take the form of partial differential equations (PDEs) to account for spatio-temporal variations in nutrient transport, cell movement, and diffusion processes.

In the context hPSC proliferation dynamics, several models have been proposed in the literature, each with unique contributions and limitations. Galvanauskas *et al*. developed a model focusing on optimizing aggregate size through controlled break-up mechanisms [27]. While effective in predicting the impact of aggregation on cell expansion, aggregate break-up was considered as a discrete intervention without mechanistically accounting for hydrodynamic forces such as shear stress and energy dissipation rate (EDR), which have been shown to play a crucial role in regulating aggregate size, cell viability, and differentiation potential in suspension cultures [28, 29]. Separately, Manstein *et al*. presented a metabolic model that predicts growth kinetics based on glucose, lactate, and glutamine dynamics, pH, and osmolality in stirred tank bioreactors (STBRs), achieving high cell densities through perfusion feeding strategies [30]. However, their model similarly lacks explicit mechanistic treatment of the hydrodynamic environment and its effects on proliferation and aggregation, and was developed for STBR systems. With the growing interest in scalable 3D culture systems, vertical wheel bioreactors (VWBRs) have emerged as an attractive platform for hPSC expansion, combining radial and axial flow components to produce more uniform distributions of hydrodynamic forces and lower shear stress compared to traditional STBRs, [31, 32, 33, 7, 34]. Studies have demonstrated that VWBRs can maintain aggregate formation within controlled size ranges during hPSC culture [31, 32, 35], and this control is critical because larger aggregates can develop necrotic cores due to limited oxygen and nutrient diffusion, adversely affecting cell growth, differentiation potential, and overall process yield [36, 37].

To address these limitations, this work develops and validates a mechanistic *in silico* model that incorporates the effects of shear stress and energy dissipation rate on hPSC proliferation and aggregate growth dynamics in a VWBR system.

## 2 Methodology

### 2.1 Experimental Setup

hPSCs were cultured as described previously [38]. For 2D static culture, undifferentiated ESI-017 human embryonic stem cells [39] (BioTime) were cultured on hESC-qualified Matrigel (Corning) coated tissue culture plastic in mTeSR1 medium (StemCell Technologies). Cells were thawed from a master cell bank and seeded at 18,000 cells/cm^2^ in mTeSR1 medium supplemented with the Rho-associated protein kinase (ROCK) inhibitor Y-27632 (10 µM, StemCell Technologies). Twenty-four hours after seeding, daily media exchanges were performed with mTeSR1 without Y-27632. Three days later, cells were passaged as single cells using Accutase (Millipore-Sigma) and seeded into Matrigel-coated 10 cm dishes (TPP Techno Plastic Products AG) using the same density as above.

For 3D hPSC culture, 100 mL and 500 mL vertical wheel bioreactors (VWBR; PBS Biotech) were used. After one passage in 2D, hPSCs were dissociated using Accutase (Millipore-Sigma) and single cells were seeded into VWBRs at the indicated cell densities in mTeSR1 medium supplemented with 10 µM Y-27632. VWBRs were operated at full volume (100 mL and 500 mL) with agitation rates stated in revolutions per minute (RPM). No media exchange was performed on day 1 after initiating the 3D culture. For condition 1 (2 × 10^4^ cells/mL starting density), a single 50% media exchange was performed on day 4 by allowing the hPSC aggregates to gravity-settle for 5–8 minutes, aspirating the spent medium, and adding fresh medium. For conditions 2–6, the medium was exchanged daily from day 2 until cell harvest. On day 2, a 50% media exchange was performed as described above. From days 3 until harvest, the 20%, 50%, or 80% of the medium was exchanged as indicated for each figure. To monitor cell density, aggregate diameter, and glucose/lactate levels, a 1 mL sample was removed from each VWBR daily, immediately prior to media exchanges at the indicated time points. Aggregates were dissociated using Accutase, after which total cell number and viability (Acridine orange/propidium iodide) was measured using a Nexcelom Cellometer K2 automated fluorescent cell counter. Aggregate diameter each day was measured manually using ImageJ (≥ 80 aggregates per replicate). On the final day of 3D culture, cells from 100 mL and 500 mL VWBRs were collected into 50 mL conical tubes, pelleted at 400 rpm for 3 minutes and dissociated with 10 mL Accutase supplemented with DNase I (∼20 U/mL; Millipore-Sigma) for 10 minutes at 37°C, pipetting the mixture once at the halfway point of the dissociation. Following enzymatic dispersion, cells were washed with 15 mL of DMEM/F12 media supplemented with DNase I and centrifuged at 1200 rpm for 5 minutes. Cell pellets were resuspended in DMEM/F12 medium, filtered through a 70 µm filter, and viable cells were quantified by using the automated cell counter described above. Media glucose and lactate levels were measured by the Laboratory Medicine Program at Toronto General Hospital (Toronto, Ontario, Canada).

### 2.2 Model Development

#### 2.2.1 Model Proposals

The mathematical framework developed in this study applies the methodology of Mostofinejad *et al*. [40] to the ODE system proposed by Galvanauskas *et al*. [27], modified to incorporate the mechanical environment in vertical-wheel bioreactor (VWBR) systems through shear stress and energy dissipation rate dependencies. The model captures the evolution of four key state variables: cell density (*X*, cells/mL), aggregate size (*Agg, µ*m), glucose concentration (*G*, mM), and lactate concentration (*Lac*, mM). Seven distinct model variants were developed to test specific hypotheses about the functional forms governing cellular responses to these hydrodynamic stimuli, drawing on established principles from biochemical kinetics and fluid mechanics.

The baseline model (Model 0) describes hiPSC expansion in a suspension culture with periodic medium exchange:

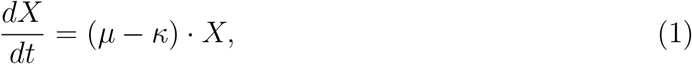

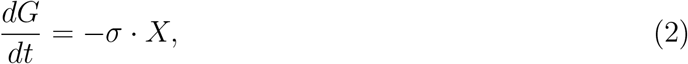

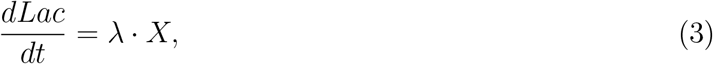

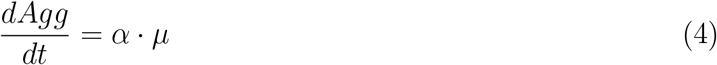

The specific growth rate *µ* follows Monod-type kinetics with multiple inhibition terms that reflect the complex metabolic constraints operating in stem cell cultures [41, 42]:

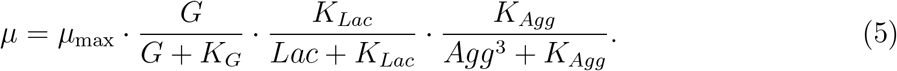

This formulation incorporates substrate limitation through the Monod term for glucose, product inhibition by lactate through a Haldane term, and diffusion limitations within cell aggregates represented by the cubic dependence on aggregate size. The cubic relationship reflects the volumetric scaling of diffusion limitations as aggregates increase in size [27].

A critical aspect of the model is the time-varying death rate that captures the initial adaptation phase of the cells to the bioreactor environment:

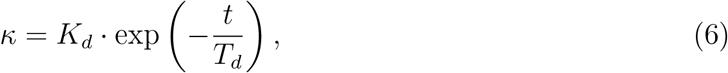

where *K*_*d*_ represents the initial death rate constant and *T*_*d*_ defines the characteristic adaptation time. This exponential decay models the death rate due to initial adaptation stress, with studies documenting this behavior during initial 24-hour expansion [43, 44, 45]. The formulation yields a maximal death rate equal to *K*_*d*_ at the beginning of the expansion process, then decreases exponentially over time [27].

The metabolic rates governing substrate consumption and product formation are coupled to growth through conversion yields. Glucose consumption accounts for glucose utilization for cell growth and lactic acid production due to overflow metabolism:

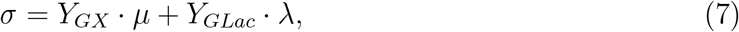

where the first term represents glucose utilization for cell synthesis linked through the glucose-to-cell conversion yield *Y*_*GX*_, and the second accounts for glucose consumed in lactate production linked through the glucose-to-lactate conversion yield *Y*_*GLac*_.

Lactate production uses a simple linear relationship to reflect the influence of elevated growth rate and glucose concentration levels on lactic acid production:

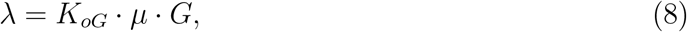

where *K*_*oG*_ is a proportionality coefficient reflecting the intensity of lactic acid production due to overflow metabolism [27].

In the extended models, the specific growth rate is modified to include shear stress effects:

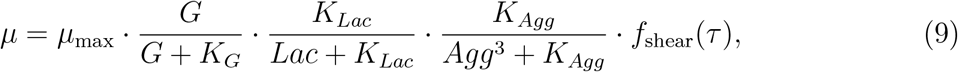

where *f*_shear_(*τ*) represents a dimensionless function describing the inhibitory effects of shear stress on cellular growth processes [26, 46, 47, 48, 49, 50, 51, 52]. The aggregate evolution equation includes a detachment term *f*_EDR_(*ε, Agg*) that models the physical breakup of cell aggregates under turbulent conditions [29, 53].

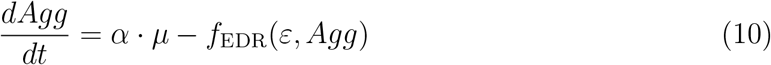

CFD simulations of the 100 mL VWBR, reported by Bauer *et al*. [54], show that shear stress and energy dissipation rate are spatially heterogeneous and concentrated near the wheel region, with both quantities increasing substantially with agitation rate (Figure 1). Volume-averaged values from these simulations were used as inputs *τ* and *ε* to the model variants.

**Figure 1:**
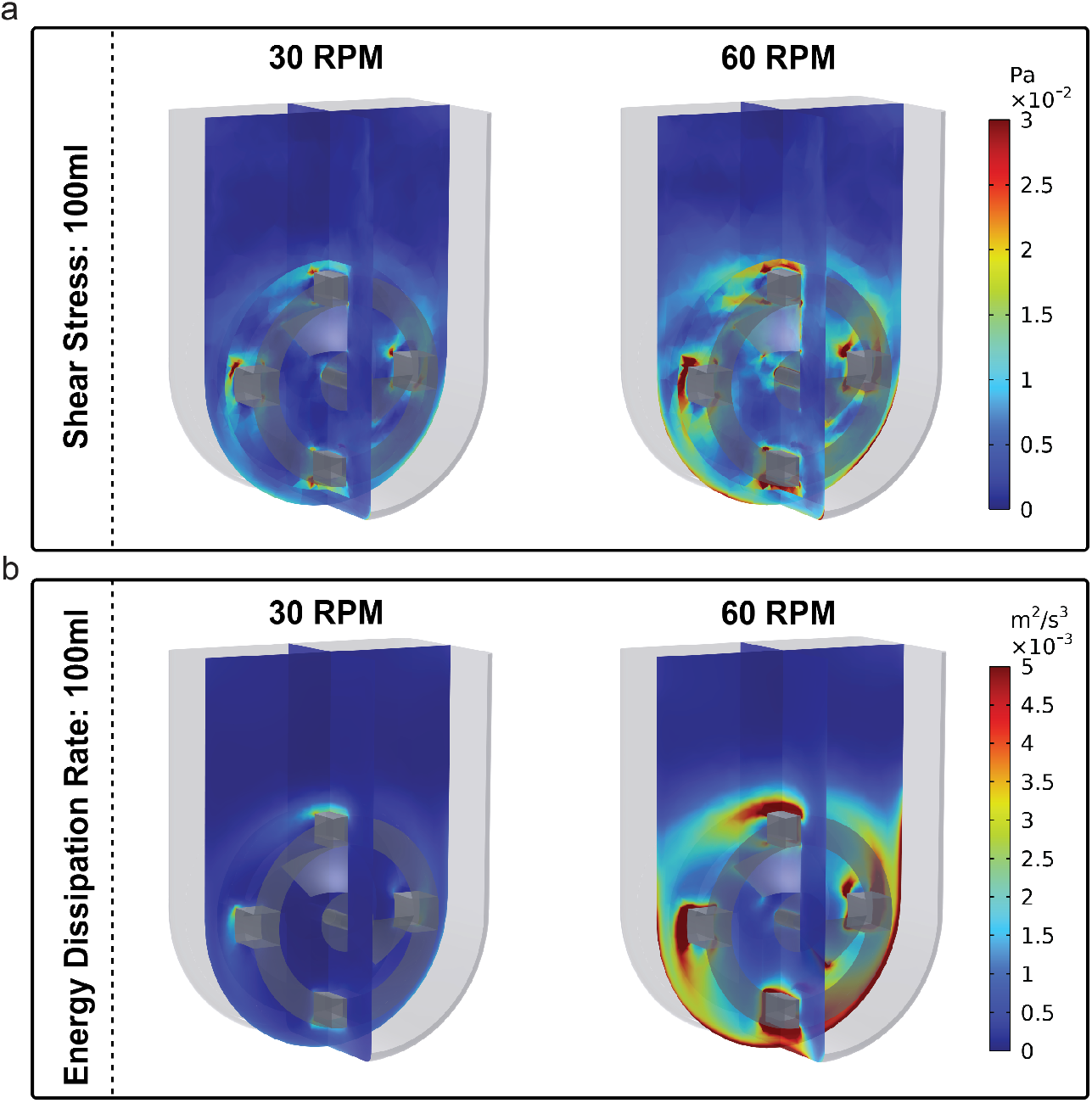
CFD-derived hydrodynamic environment in the 100 mL VWBR at 30 and 60 RPM. (a) Shear stress (Pa) and (b) energy dissipation rate (m^2^ s^-3^) distributions on a mid-plane cross-section. Elevated values concentrate near the wheel regions and increase with agitation rate. Adapted from Bauer *et al*. [54].

The seven model variants differ in their functional forms for *f*_shear_(*τ*) and *f*_EDR_(*ε, Agg*), testing specific hypotheses about cellular responses to hydrodynamic stresses (Table 1).

**Table 1:** Model variants testing different functional forms for shear stress and energy dissipation rate (EDR) effects. *K*_*τ*_ = shear half-saturation constant; *K*_*ε*_ = EDR half-saturation constant; *δ*_max_ = maximum detachment rate.

| Model | Name | $f_{\text{shear}}(\tau)$ | $f_{\text{EDR}}(\varepsilon, \text{Agg})$ | Hypothesis |
| --- | --- | --- | --- | --- |
| 1 | Michaelis-Menten | $\frac{K_\tau}{\tau + K_\tau}$ | $\delta_{\max} \cdot \frac{\varepsilon}{\varepsilon + K_\varepsilon} \cdot \text{Agg}$ | Adaptive cellular mechanisms become overwhelmed at high shear stress [55, 46, 23, 56, 57, 58, 59, 60, 61] |
| 2 | Linear Competitive | $\frac{1}{1 + \tau/K_\tau}$ | Same as Model 1 | Shear acts as competitive inhibitor without beneficial effects [42] |
| 3 | Exponential Approximation | $\frac{1}{1 + x + x^2/2}$<sup>a</sup> | Same as Model 1 | Exponential decay behavior via Taylor expansion [62] |
| 4 | Growth-Only Shear | Same as Model 1 | Same as Model 1 | Shear affects growth while leaving metabolic processes unaffected [51]<sup>b</sup> |
| 5 | Hill Shear (n=2) | $\frac{K_\tau^2}{\tau^2 + K_\tau^2}$ | Same as Model 1 | Sharp threshold response via cooperative inhibition [63] |
| 6 | Growth-Coupled EDR | Same as Model 1 | $\mu \cdot \frac{\varepsilon}{K_\varepsilon} \cdot \text{Agg}$ | Aggregate breakup is coupled to growth, so detachment ceases when proliferation stalls<sup>d</sup> |
| 7 | Dual Interaction | $\frac{K_{\tau,\text{eff}}}{\tau + K_{\tau,\text{eff}}}$<sup>c</sup> | Same as Model 1 | Turbulent fluctuations modify cellular sensitivity to shear stress [64, 65, 66] |

<sup>a</sup> $x = \tau/K_\tau$ .
<sup>b</sup> Metabolic rates use $\mu_{\text{base}}$ instead of $\mu_{\text{eff}}$ .
<sup>c</sup> $K_{\tau,\text{eff}} = K_\tau \cdot (1 + \gamma \cdot \varepsilon/(\varepsilon + K_\varepsilon))$ , where $\gamma$ is the interaction parameter.
<sup>d</sup> For Model 6, $f_{\text{EDR}}$ also depends on $\mu$ ; the aggregate equation reads $d\text{Agg}/dt = \mu(\alpha - \varepsilon/K_\varepsilon \cdot \text{Agg})$ .

#### 2.2.2 Structural Identifiability Analysis

Structural identifiability analysis determines whether model parameters can be uniquely determined from experimental data, assuming noise-free measurements. To assess structural identifiability, the models are expressed as nonlinear dynamical systems:

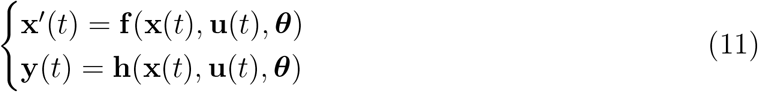

where **x**(*t*) = [*X*(*t*), *G*(*t*), *Lac*(*t*), *Agg*(*t*)]^*T*^ is the time-dependent state vector containing cell density, glucose concentration, lactate concentration and aggregate size, **u**(*t*) = [*τ* (*t*), *ε*(*t*)]^*T*^ denotes the input vector of shear stress and energy dissipation rate, *θ* represents the set of unknown parameters, and **y**(*t*) represents the outputs which are cell density, aggregate size, glucose concentration, and lactate concentration with **y**(*t*) = **x**(*t*).

Identifiability analysis employs differential algebra and symbolic computation methods to construct the observability matrix **O**(*θ*) of the non-linear dynamical system through successive derivatives of the output equations. A model is structurally identifiable if rank(**O**(*θ*)) = dim(*θ*).

#### 2.2.3 Objective Function Definition

Parameter estimation requires defining an objective function that quantifies the goodness-of-fit between model predictions and experimental data. Two standard methods for parameter estimation are maximum likelihood estimation (MLE) and nonlinear least squares (NLS) [15, 67]. MLE estimates parameters by maximizing the likelihood function, which represents the probability of observing the experimental data given a set of model parameters. In this work, MLE is employed because it provides a suitable probabilistic framework for the parameter estimation process and is consistent with identifiability analysis methods that rely on profile likelihood approaches.

Assuming that experimental noise follows a normal distribution, the likelihood function is expressed as:

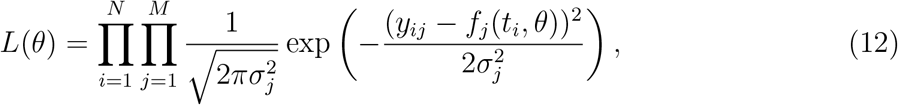

where *θ* represents the set of model parameters, *y*_*ij*_ is the experimental observation for observable *j* at time *t*_*i*_, *f*_*j*_(*t*_*i*_, *θ*) is the model prediction, 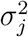 is the variance of observable *j*, and *N* and *M* denote the number of time points and observables, respectively. Taking the negative log-likelihood yields:

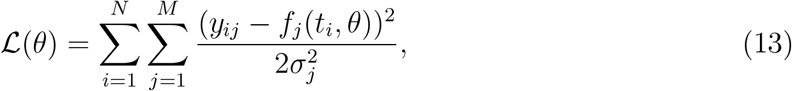

which is mathematically equivalent to the weighted NLS formulation under the assumption of normally distributed errors.

#### 2.2.4 Model Calibration and Selection Using Experimental Data

Following the acquisition of experimental data, the data is randomly split into three subsets for calibration (60%), selection (20%), and validation (20%) [40]. The calibration data set is then used to formulate a nonlinear optimization problem for each candidate model to determine the set of model parameters that best fit the data. This involves minimizing the objective function ℒ (*θ*), which quantifies the discrepancy between model predictions and experimental observations, namely the negative log-likelihood function of Eq. 13. The optimization problem is expressed as:

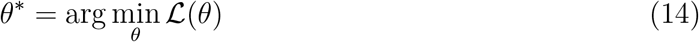

where *θ* represents the set of model parameters to be inferred, and *θ*^∗^ is the optimal parameter set that minimizes ℒ (*θ*).

To systematically explore the parameter space and avoid local minima, a max-min Latin hypercube sampling (LHS) method is utilized to generate a diverse set of initial parameter guesses [40, 68, 69]. Using each initial guess as a starting point, the optimization problem is first solved using the Black Box Optimization (BBO) algorithm in Julia, which efficiently handles non-linear, non-convex functions without requiring gradient information [70]. The best-performing solutions from this stage are then used as starting points for further optimization with the Broyden-Fletcher-Goldfarb-Shanno (BFGS) method, a quasi-Newton algorithm that leverages gradient information to ensure final convergence to an optimal solution [71, 72].

After calibration, model selection is made between the candidate models by computing the Akaike Information Criterion (AIC) and Bayesian Information Criterion (BIC) for the selection dataset, namely:

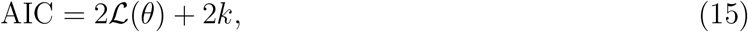

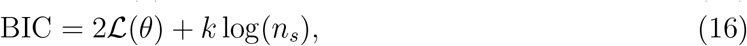

where *k* is the number of inferred parameters, and *n*_*s*_ is the number of observations in the selection dataset. The model with the lowest AIC or BIC value is selected as the best descriptor for the experimental data, balancing model complexity and goodness of fit.

#### 2.2.5 Practical Identifiability Analysis

Practical identifiability analysis is performed to assess whether the inferred model parameters can be reliably determined given the available, noisy experimental data. [19].

To evaluate practical identifiability, the profile likelihood approach is employed, which examines how variations in an individual parameter affect the objective function while allowing all other parameters to re-optimize [73]. The profile likelihood for a given parameter *θ*_*k*_ is defined as:

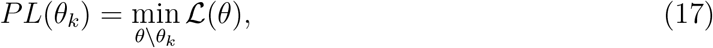

where ℒ (*θ*) is the objective function, *θ* = (*θ*_1_, *θ*_2_, …, *θ*_*m*_) is the full set of model parameters, *θ*_*k*_ is the parameter being profiled, and *θ \θ*_*k*_ represents all parameters except *θ*_*k*_, which are optimized to minimize ℒ (*θ*) at fixed values of *θ*_*k*_.

A well-identified parameter exhibits a sharply peaked profile likelihood curve, meaning small deviations from the optimal 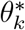 significantly increase the objective function. Conversely, a flat or broad likelihood profile suggests weak parameter sensitivity, leading to high uncertainty in parameter estimation. Confidence intervals for *θ*_*k*_ can be derived from the profile likelihood using the threshold:

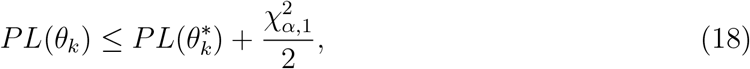

where 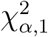 is the chi-square quantile corresponding to the desired confidence level 1 − *α*.

#### 2.2.6 Goodness of Fit

Performance of the selected model is assessed using the Mean Relative Prediction Error (MRPE) as the primary goodness-of-fit metric. MRPE quantifies the average relative difference between model predictions and experimental measurements, providing a normalized measure of model error that allows comparison across different types of observables. This metric is particularly useful in biological systems, where measurements can span multiple orders of magnitude.

The MRPE is computed as:

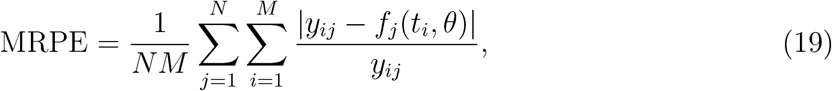

where *N* is the number of time points, *M* is the number of observables, *y*_*ij*_ is the experimental observation for observable *j* at time *t*_*i*_ and, *f*_*j*_(*t*_*i*_, *θ*) is the model prediction for the same observable and time point. A lower MRPE value indicates a better fit, as it reflects smaller relative discrepancies between model predictions and experimental data.

To contextualize model performance, the mean relative experimental variance of the experimental data was calculated. This metric quantifies the inherent variability in the experimental measurements. Specifically, the mean relative experimental variance is calculated as the standard deviation among the number of experiment replicates of each observable at each time point, normalized with the value of the mean value of the observable at that time point, and averaged over all time points and observables:

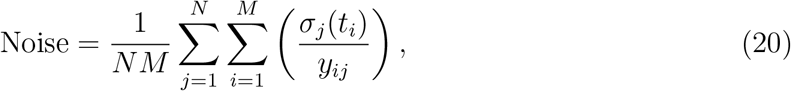

where *σ*_*j*_(*t*_*i*_) is the standard deviation of replicate measurements for observable *j* at time point *i*.

Comparing the model MRPE with the mean relative experimental variance provides valuable context for model evaluation. When the MRPE approaches the experimental noise, it suggests that the model captures the underlying biological processes to the extent permitted by measurement error.

## 3 Results

### 3.1 Model Development

#### 3.1.1 Structural Identifiability Analysis

The baseline model from Galvanauskas et al. [27] was initially examined for structural identifiability and found to be not fully identifiable. Specifically, the death rate, expressed as *K*_*d*_ × exp(−*t/T*_*d*_), where both *K*_*d*_ and *T*_*d*_ needed to be estimated from experimental data, was not structurally identifiable. This exponential formulation creates structural identifiability problems because multiple combinations of (*K*_*d*_, *T*_*d*_) parameters can produce identical model outputs. To address this issue, in this work we reformulated the death rate using a single identifiable parameter *κ* = *K*_*d*_ × exp(−*t/T*_*d*_) with *T*_*d*_ fixed at 12 hours. This value was selected from the range of 6-24 hours reported in stem cell culture literature for characteristic times of cellular adaptation and stress response in media environments [43, 44, 45]. This modified model was verified to be structurally identifiable.

Structural identifiability analysis was performed on the other seven candidate models presented in Sec. 2.2.1 using StructuralIdentifiability.jl, an open-source SciML package [74, 75, 76]. The analysis revealed that all seven candidate models (Models 1–7) achieved complete global identifiability.

#### 2.2.4 Model Calibration and Selection Using Experimental Data

All eight models were calibrated using experimental data from six different bioreactor conditions, including variations in initial cell density (2 × 10^4^, 8 × 10^4^, 15 × 10^4^ cells/mL), agitation rate (30, 40, 60 RPM), and working volume (100 mL, 500 mL). A summary of the experimental conditions used in this study is shown in Table 2. Conditions 1, 2, 3, and 6 were conducted over 6 days, while conditions 4 and 5 were conducted over 5 days.

**Table 2:** Experimental conditions used to generate data for model calibration, selection and validation.

| Exp # | Initial Cell Density [cells/mL] | Agitation Rate [rpm] | Bioreactor Volume [mL] |
| --- | --- | --- | --- |
| 1 | $2 \times 10^4$ | 40 | 100 |
| 2 | $8 \times 10^4$ | 40 | 100 |
| 3 | $15 \times 10^4$ | 40 | 100 |
| 4 | $8 \times 10^4$ | 30 | 500 |
| 5 | $8 \times 10^4$ | 30 | 100 |
| 6 | $8 \times 10^4$ | 60 | 100 |

The calibration process employed a multi-start optimization strategy as described in the methods section. After all parameters for the candidate models were inferred with the calibration set, the selection dataset was used to calculate BIC values for all models. BIC results, shown in Figure 2, revealed that Model 6 achieved the lowest BIC value and was therefore selected as the best-performing model.

**Figure 2:**
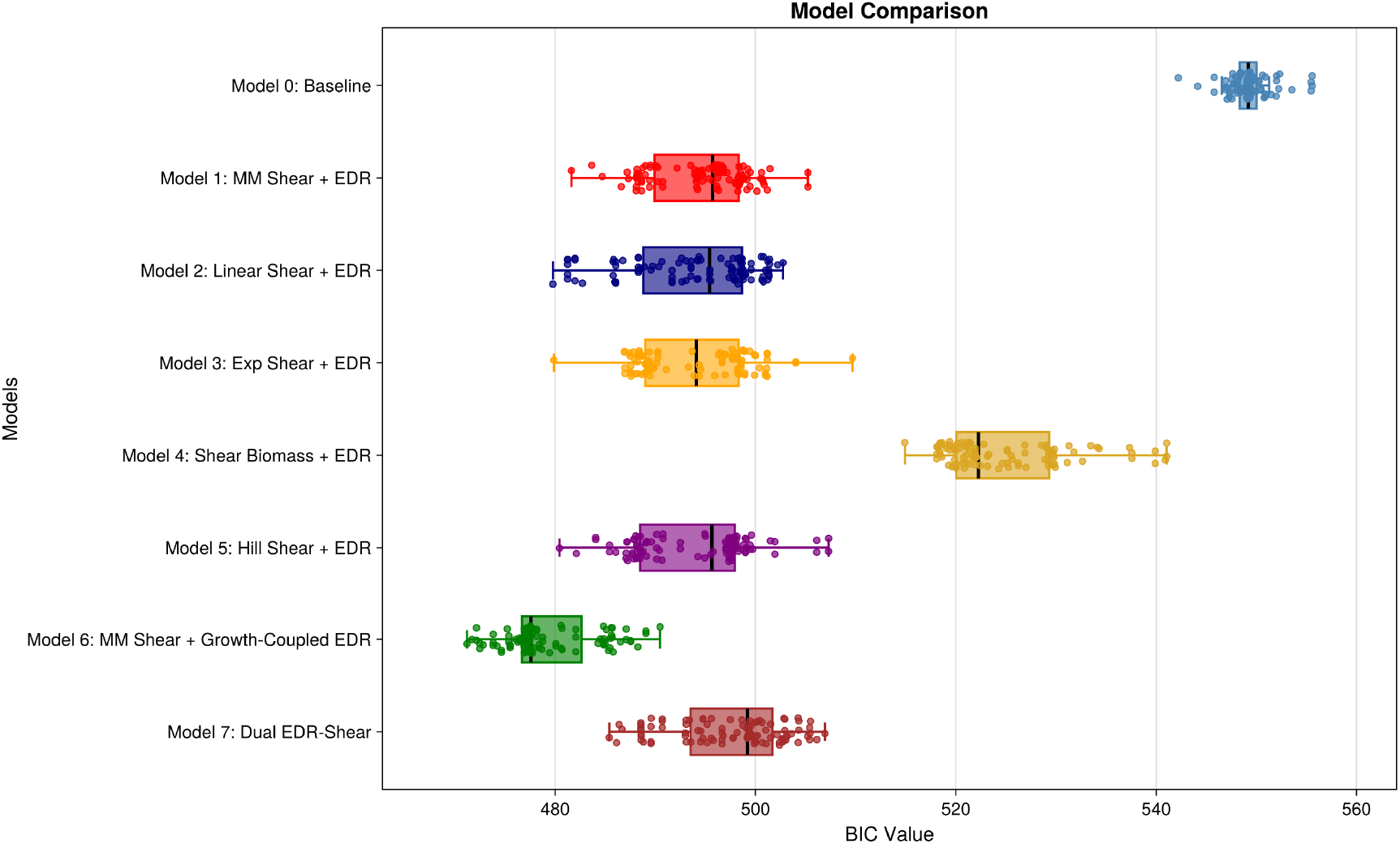
Bayesian Information Criterion analysis for model selection. BIC values for all eight model variants, with Model 6 achieving the lowest BIC value and thus selected as the best-performing model.

Table 3 presents the inferred parameter values for Model 6 with their corresponding confidence intervals and units. The parameter estimation procedure successfully determined values for all 11 model parameters, including the baseline model parameters and the parameters for shear stress and energy dissipation rate effects.

**Table 3:** Inferred parameter values for Model 6 with confidence intervals and units using experimental data.

| Parameter | Value | (Lower Bound, Upper Bound) | Units |
| --- | --- | --- | --- |
| $K_{\text{Agg}}$ | $6.98 \times 10^7$ | $(3.34 \times 10^7, 4.38 \times 10^8)$ | $\mu\text{m}^3$ |
| $K_d$ | 0.133 | (0.110, 0.151) | $\text{h}^{-1}$ |
| $K_G$ | 14.59 | (2.17, 30.0) | mM |
| $K_{\text{Lac}}$ | 34.75 | Fixed<sup>a</sup> | mM |
| $K_{\text{oG}}$ | $3.20 \times 10^{-6}$ | $(2.75 \times 10^{-6}, 3.24 \times 10^{-6})$ | mL/cell |
| $Y_{\text{GLac}}$ | 0.573 | (0.496, 0.567) | mmol/mmol |
| $Y_{\text{GX}}$ | $9.43 \times 10^{-7}$ | Fixed<sup>a</sup> | mmol/cell |
| $\alpha$ | 71.79 | (60.4, 73.5) | $\mu\text{m}$ |
| $\mu_{\text{max}}$ | 0.133 | (0.0681, 0.205) | $\text{h}^{-1}$ |
| $K_\tau$ | $8.32 \times 10^{-3}$ | $(5.51 \times 10^{-3}, 1.00 \times 10^{-2})$ | Pa |
| $K_\varepsilon$ | $4.78 \times 10^{-3}$ | $(4.28 \times 10^{-3}, 9.52 \times 10^{-3})$ | $\text{m}^2 \text{ s}^{-3}$ |

<sup>a</sup> Fixed at maximum likelihood estimate from initial calibration; confidence interval not applicable.

The inferred Model 6 demonstrates really good predictive capability across all experimental conditions, as shown in Figure 3. The model successfully captures the temporal dynamics of cell density, aggregate size, glucose consumption, and lactate production under diverse biochemical and fluid dynamic conditions. Cell density predictions show good agreement with experimental trends, successfully reproducing the growth patterns observed under different initial seeding densities and agitation conditions. Aggregate size predictions capture the general trends of aggregate formation dynamics under varying fluid dynamic conditions, though the model shows some over- and under-prediction relative to the experimental data. For glucose and lactate concentrations, the model accurately represents both consumption and production patterns, including the effects of media changes evident in the experimental data. Notably, the model predictions consistently fall within the experimental error bars across most time points, indicating strong agreement between *in silico* results and *in vitro* observations. The model shows balanced performance in both under- and over-predicting the measured state variables, with deviations typically remaining within the natural variability of the experimental data as defined by the standard deviation across replicates.

**Figure 3:**
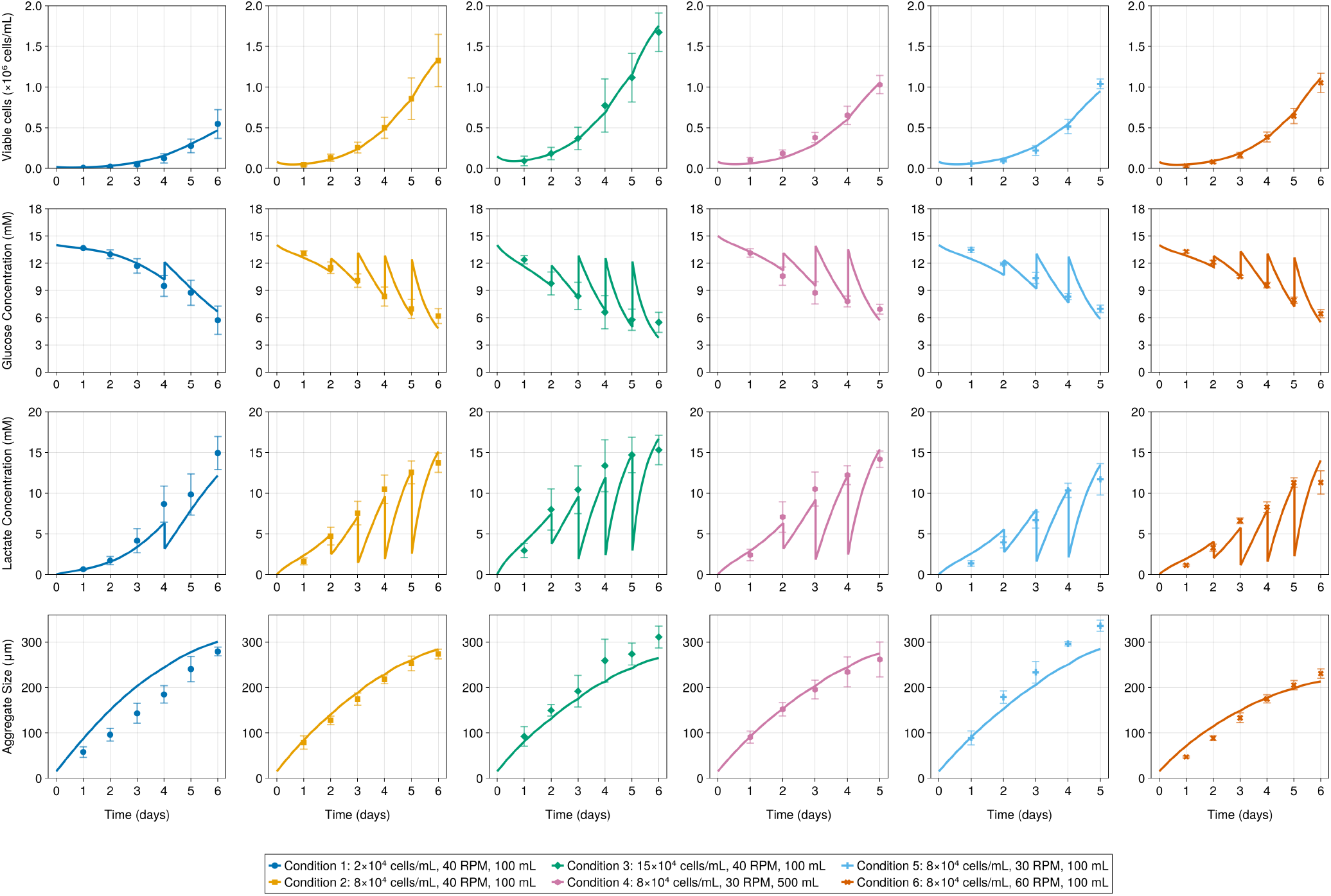
Inferred *in silico* Model 6 versus *in vitro* experimental data for cell density, aggregate size, glucose concentration, and lactate concentration (columns) across six bioreactor conditions (rows). Solid lines represent Model 6 predictions, while experimental data points are shown with solid markers. Different colors represent the six experimental conditions 1 to 6. Error bars indicate standard deviation across replicates (*n* = 3 for each condition). Conditions 1, 2, 3, and 6 span 6 days while Conditions 4 and 5 span 5 days. Peaks observed in the glucose and lactate concentration plots correspond to scheduled media changes, where fresh media replenishes glucose levels while simultaneously removing or diluting accumulated lactate.

#### 3.1.3 Practical Identifiability Analysis

Practical identifiability of the selected model was assessed through profile likelihood analysis using the `ProfileLikelihood.jl` package [77]. Profile likelihood confidence intervals were computed by fixing each parameter in turn across a defined range and optimizing the remaining parameters to obtain the profile log-likelihood, 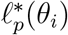.

An initial 11-parameter analysis revealed that two parameters (*K*_Lac_ and *Y*_*GX*_) were practically non-identifiable from the available experimental data. The lactate inhibition constant *K*_Lac_ was non-identifiable from above: because lactate concentrations remained well below inhibitory levels throughout all experimental conditions, the term *K*_Lac_*/*([Lac]+*K*_Lac_) ≈ 1 at all times, rendering *K*_Lac_ structurally insensitive to the data. The biomass yield coefficient *Y*_*GX*_ was non-identifiable from below: glucose consumption was dominated by the overflow (lactate) pathway governed by *Y*_GLac_, making the direct oxidative yield *Y*_*GX*_ a negligible contributor to the overall glucose balance. These two parameters were therefore fixed at their maximum likelihood estimates from the initial calibration. Notably, the fixed value of *K*_Lac_ is of the same order of magnitude as published estimates for hPSC culture, including the range of 45–65 mM reported by Galvanauskas *et al*. [27] and Manstein *et al*. [30], lending some support to this choice. The profile likelihood analysis was then repeated for the remaining nine free parameters.

Figure 4 shows the resulting profiles for all nine free parameters. All profiles are smooth, unimodal, and cross the 95% confidence threshold at finite values on both sides, confirming that the model is practically identifiable. The corresponding parameter values and 95% confidence intervals are reported in Table 3.

**Figure 4:**
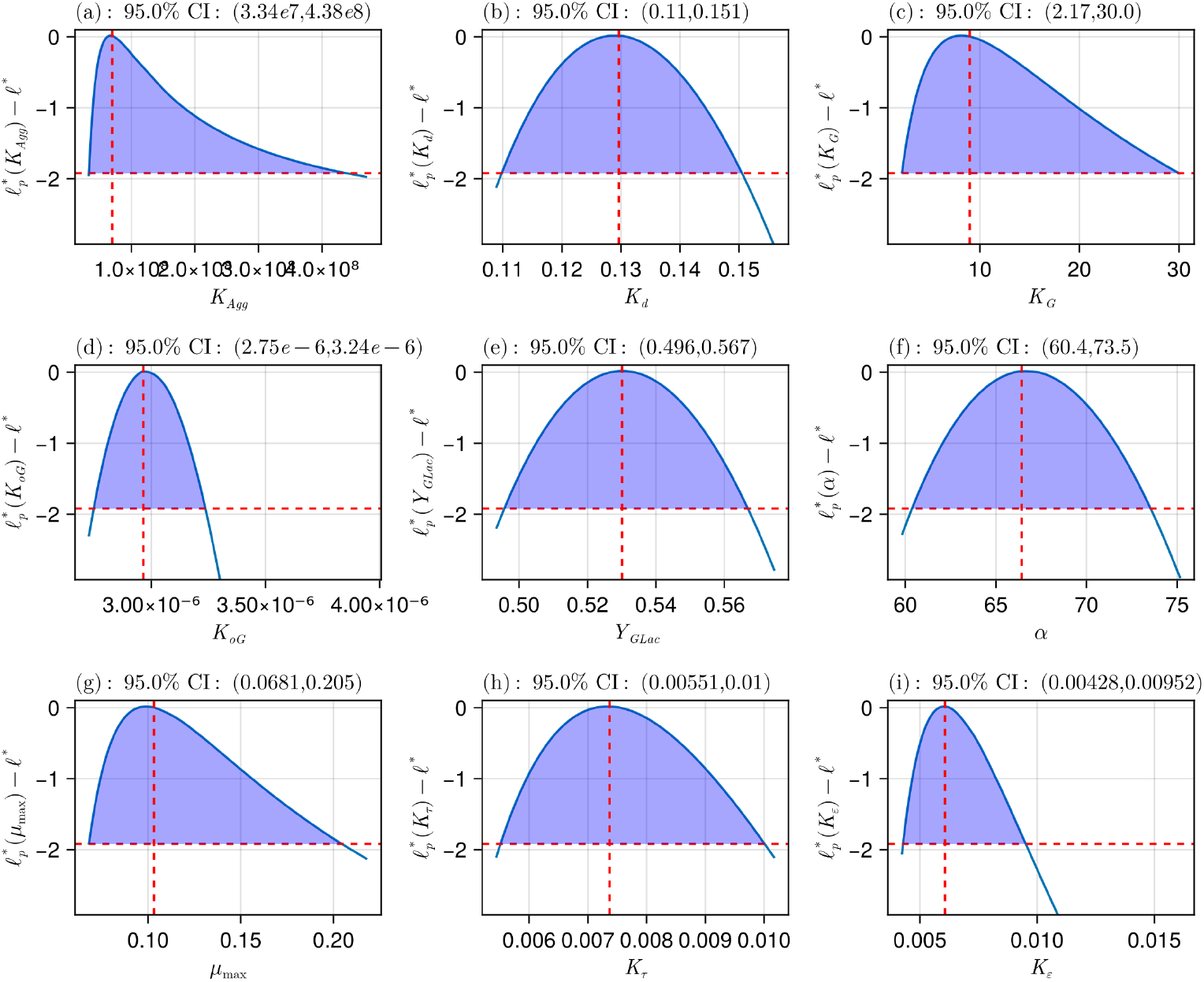
Profile likelihood analysis for model parameters. Each subplot shows the profile likelihood as a function of the parameter value. The red vertical line indicates the inferred parameter value, while the red horizontal line marks the 95% confidence threshold. The intersections of the profile curves with the horizontal threshold line define the 95% confidence intervals for each parameter, demonstrating practical identifiability of all model parameters.

#### 3.1.4 Goodness of Fit

The MRPE of the inferred model was calculated as described in the methodology and found to be 13.97%. To assess the accuracy of the model relative to experimental variability, the experimental noise, defined as the mean relative standard deviation of experimental replicates across all time points and conditions, was computed and found to be 16.29%. The close alignment between these values indicates that the model’s predictive error is comparable to the inherent variability in the experimental data, demonstrating that the inferred model provides a sufficiently accurate representation of the observed biological system.

## 4 Model Application: Protocol Optimization

Having established Model 6 through calibration across multiple experimental conditions, the model was applied to investigate the influence of media exchange percentage on process yield. A parametric study was conducted using the calibrated model, with predictions subsequently confirmed through targeted *in vitro* experiments using the same ESI017 embryonic stem cell line. The percentage of media exchanged (20%, 50%, and 80% baseline) was varied to determine its influence on cell expansion and metabolite dynamics. All experiments used initial cell density, agitation rate, and working volume corresponding to Condition 2 (Table 2), following the protocol described in Sec. 2.1.

Figure 5 compares model predictions with experimental measurements for cell density, glucose, lactate, and aggregate size across the 6-day culture period. The model demonstrates strong predictive accuracy, with MRPEs of 12.16% and 13.91% for the P20 and P50 conditions, respectively, comparable to experimental noise levels (12.09% and 9.84%), indicating predictions fall within experimental variability, although the model slightly underpredicts cell population. Model predictions show final cell densities of 1.30 × 10^6^, 1.13 × 10^6^, and 0.93 × 10^6^ cells/mL for P80, P50, and P20, representing 13.5% and 28.7% reductions from baseline, respectively. The P50 condition maintains higher net glucose availability in early culture due to greater replenishment per exchange, sustaining faster growth. However, the resulting larger cell population drives greater glucose consumption and lactate accumulation in later days, such that P50 exhibits lower final glucose and higher final lactate concentrations than P20. The P20 condition, while producing fewer cells, maintains a less metabolically stressed environment throughout the culture period. Aggregate size remains similar across conditions (∼264–282 *µ*m by day 6), indicating media exchange percentage primarily affects proliferation rather than aggregate formation. These results suggest P50 provides a practical balance between cell expansion and resource efficiency.

**Figure 5:**
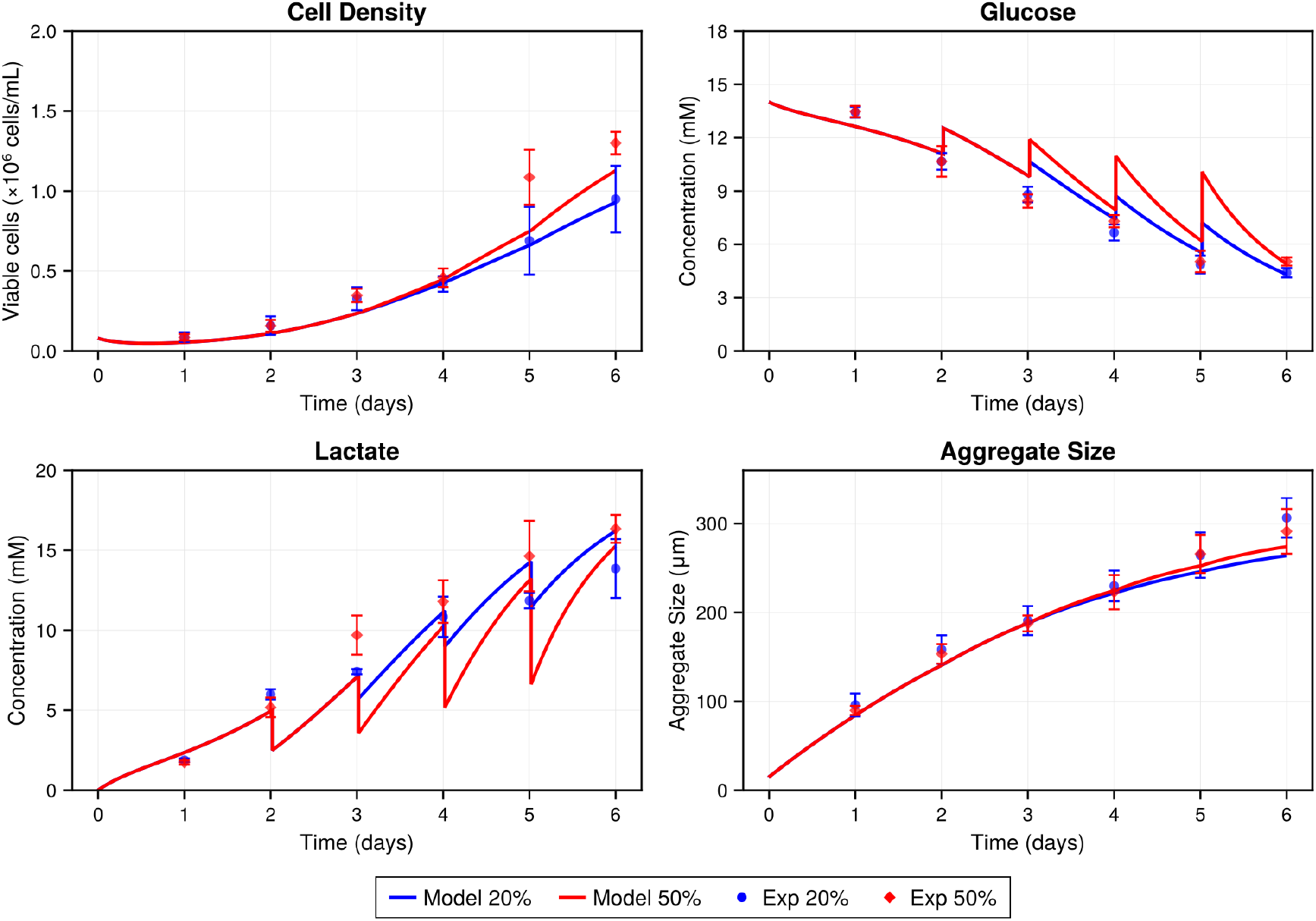
Model application to media exchange protocol optimization. Comparison of model predictions (solid lines) with experimental data (markers with error bars) for 20% (blue) and 50% (red) media exchange using ESI017 embryonic stem cells at 8 × 10^4^ cells/mL initial density and 40 RPM. The four panels show (top left) viable cell density, (top right) glucose concentration, (bottom left) lactate concentration, and (bottom right) aggregate size over 6 days. Error bars represent standard deviation across experimental replicates (*n* = 3).

## 5 Discussion

This study presents a validated mechanistic *in silico* model that couples the hydrodynamic environment of a vertical-wheel bioreactor to hPSC proliferation and aggregate dynamics, supporting prospective prediction across operating conditions not used during calibration.

Model 6 performed best under BIC penalization, and its structure reflects the distinct mechanisms through which shear stress and energy dissipation rate act on cells and aggregates. The Michaelis-Menten form for shear stress inhibition captures a gradual, saturable impairment of cell growth consistent with known mechanosensory biology: under moderate shear, hPSCs activate protective responses including cytoskeletal reorganization and focal adhesion remodeling [56, 58, 59], which become progressively overwhelmed at higher stress levels [60, 61]. The inferred *K*_*τ*_ = 8.32 × 10^−3^ Pa places the operating window in a regime of 31–48% shear-mediated growth inhibition, confirming that shear stress is neither negligible nor fully saturating across experimental conditions [11, 50, 48, 51, 52]. The poor performance of Model 4, which assumed shear stress affects growth alone while leaving metabolic processes unaffected, further supports the view that mechanical stress broadly alters cellular physiology, including glucose consumption and lactate production [52, 51].

The distinct treatment of energy dissipation rate in Model 6 further reinforces this mechanistic picture. The growth-coupled EDR formulation implies that actively proliferating aggregates are more susceptible to fragmentation, consistent with transient weakening of intercellular junctions during cytokinesis [29, 53]. Coupling detachment to *µ* provides an intrinsic feedback whereby fragmentation diminishes as growth slows under nutrient limitation, addressing the limitation of the discrete breakup intervention used by Galvanauskas *et al*. [27].

Compared to prior hPSC proliferation models, the present framework represents a meaningful advance in mechanistic scope. Galvanauskas *et al*. [27] implemented aggregate breakup as a discrete chemical intervention using botulinum hemagglutinin at a fixed time point, followed by physical disruption by pipetting, with no link between fragmentation or cell growth and the hydrodynamic environment. Manstein *et al*. [30] employed a Monod-based metabolic model incorporating glucose, lactate, glutamine, osmolality, and aggregate diameter, achieving cell density prediction errors of 2–5% under tightly controlled STBR perfusion. Despite supplementing Pluronic F68 as a shear protectant, it is well established that Pluronic F68 reduces but does not eliminate shear-induced cell damage [51, 52], and their model contains no explicit shear stress or EDR terms. By embedding these effects directly into the kinetic structure of the ODE system, the present model captures the continuous interplay between fluid dynamics and cell behavior without requiring condition-specific interventions.

Building on this mechanistic foundation, the model was applied to media exchange protocol optimization, reframing the design problem from empirical trial-and-error to quantitative trade-off analysis. The disproportionate yield penalty from reducing exchange from 50% to 20%, relative to the modest penalty from 80% to 50%, reveals a nonlinear metabolic response difficult to identify without mechanistic insight. The insensitivity of aggregate size to exchange percentage further decouples these two design variables. A 37.5% reduction in media consumption with only a 13.5% reduction in cell yield represents a meaningful efficiency gain given the substantial cost of specialty media in clinical-scale hPSC production [11, 7, 14, 6]. Calibration across both 100 mL and 500 mL VWBR configurations suggests that volume-averaged CFD-derived hydrodynamic inputs are sufficient to capture dominant mechanical effects at these scales, and the mechanistic model structure is expected to remain applicable at larger volumes provided operating conditions remain within the calibrated range.

The model has several limitations that should be considered when interpreting and applying its predictions. The homogeneous population assumption does not capture heterogeneity in gene expression, metabolic activity, and mechanical properties known to exist in hPSC populations [5, 10]. Aggregate size is represented as a single characteristic length scale rather than a full population distribution, precluding explicit tracking of mass transfer gradients within larger aggregates [37, 36]. The complete-mixing assumption neglects spatial gradients in *τ* and *ε*, and coupling the ODE model to CFD-derived spatial fields would address this without sacrificing computational tractability [31]. Nevertheless, the inferred shear inhibition range of 31–48% across experimental conditions contextualizes the VWBR’s design advantage: operating in a regime where shear stress is biologically significant but not fully saturating suggests that the VWBR’s more uniform hydrodynamic environment [31, 33] enables meaningful mechanosensory signaling without crossing into damaging shear levels, a balance that would be more difficult to achieve in traditional stirred tank configurations. The model was validated using a single hESC line (ESI017) and does not track pluripotency markers or lineage commitment; recalibration would be required for other cell lines, and extension to differentiation protocols would require additional state variables [12, 11]. Despite these limitations, the framework provides a transferable foundation for quantitative bioprocess design essential for constraining the parameter space during scale-up and reducing reliance on empirical optimization in hPSC manufacturing [13, 14, 15].

## 6 Conclusion

This work establishes a validated mechanistic model for hPSC proliferation in vertical-wheel bioreactors that incorporates shear stress and energy dissipation rate effects on cell growth and aggregate dynamics. The systematic comparison of model variants revealed that a growth-coupled EDR formulation best describes aggregate detachment, while Michaelis-Menten kinetics captures cellular adaptation to shear stress. The model demonstrates strong predictive accuracy across diverse experimental conditions and provides a quantitative tool for optimizing culture protocols, contributing to the development of scalable, cost-effective bioprocesses for hPSC manufacturing.

## Abbreviations

hPSCs: human pluripotent stem cells
VWBR: vertical wheel bioreactor
EDR: energy dissipation rate.

## Author contributions

F.R.A., D.R., A.M. and C.A. developed the mathematical models and conducted all *in silico* studies, using shear stress and energy dissipation rates estimated by J.B. from CFD simulations of VWBR. F.J.A., C.M. and M.L. performed all the experimental work generating data used in this manuscript. F.R.A. and D.R. drafted the manuscript, while D.R., F.J.A., M.L. and C.A. reviewed and approved the final manuscript.

## Acknowledgments

We acknowledge the support of the Government of Canada’s New Frontiers in Research Fund (NFRF) [NFRFT-2022-00447, NFRFT-2020-00787], the Canada Research Chairs Program (CRC-2020-00245), and the Natural Sciences and Engineering Research Council of Canada (NSERC). We also thank Pouyan Vatani for conducting the CFD simulations and generating the hydrodynamic visualizations of shear stress and energy dissipation rate distributions in the vertical-wheel bioreactor.

## Conflict of interest

Michael A. Laflamme is a scientific founder and consultant for Bluerock Therapeutics LP (Boston, MA, USA). The other co-authors have no conflicts of interest to declare.

## Ethics Approval

All human pluripotent stem cell work was approved by the Stem Cell Oversight Committee of the Canadian Institutes of Health Research.

## Data Availability

Data and code used to generate the results in this manuscript will be made available on a public, archival repository (Zenodo) upon acceptance of the manuscript.

